# Elevated phosphorylation of EGFR in NSCLC due to mutations in PTPRH

**DOI:** 10.1101/2021.09.14.460311

**Authors:** Matthew R. Swiatnicki, Jonathan P. Rennhack, Daniel P. Hollern, Ashlee V. Perry, Rachel Kubiak, Sarai M. Riveria Riveria, Sandra O’Reilly, Eran R. Andrechek

## Abstract

The role of EGFR in lung cancer is well described with numerous activating mutations that result in phosphorylation and tyrosine kinase inhibitors that target EGFR. While the role of the EGFR kinase in non-small cell lung cancer (NSCLC) is appreciated, control of EGFR signaling pathways through dephosphorylation by phosphatases is not as clear. In recent work we identified mutations in Protein Tyrosine Phosphatase Receptor Type H (*Ptprh*, also known as SAP-1) as being associated with elevated phosphorylation of EGFR in a mouse model of breast cancer. We have examined a series of tumors from this mouse model, revealing conserved V483M *Ptprh* mutations within the FVB background, but a series of varied mutations in other backgrounds. Despite the varied *Ptprh* mutations in other background strains, matched primary and metastatic tumors largely shared mutational profiles. Profiling the downstream events of *Ptprh* mutant tumors revealed AKT activation, suggesting a key target of PTPRH was EGFR tyrosine 1197. Given the role of EGFR in lung cancer, we explored TCGA data which revealed that a subset of *PTPRH* mutant tumors shared gene expression profiles with *EGFR* mutant tumors, but that *EGFR* mutations and *PTPRH* mutations were mutually exclusive. Generation of a PTPRH knockout NSCLC cell line resulted in Y1197 phosphorylation of EGFR, and a rescue with expression of wild type PTPRH returned EGFR phosphorylation to parental line values while a rescue with a D986A catalytically dead mutant PTPRH did not, demonstrating that PTPRH targets EGFR. As expected with active EGFR, the knockout of PTPRH was associated with increased growth rate. Moreover, a dose response curve illustrated that two human NSCLC lines that had naturally occurring *PTPRH* mutations responded to EGFR tyrosine kinase inhibition. Injection of one of the NSCLC human lines into mice resulted in tumors, and Osimertinib treatment resulted in a reduction of tumor volume relative to vehicle controls. Consistent with prior literature from breast cancer, *PTPRH* mutation resulted in nuclear pEGFR as seen in immunohistochemistry, suggesting that there may also be a role for EGFR as a transcriptional co-factor. Other roles for PTPRH were explored through a receptor tyrosine kinase array, noting elevated phosphorylation of FGFR1. Knockout of PTPRH in NSCLC cell lines resulted in elevated phosphorylated FGFR1 relative to controls, indicating that PTPRH has a number of targets that may be aberrantly activated in NSCLC with mutations in PTPRH. Together these data suggest that mutations in PTPRH in NSCLC may result in clinically actionable alterations using existing therapies.

## Introduction

Lung cancer results in the greatest number of U.S. cancer deaths in both men and women, and 5 year survival rates remain poor^1^. Lung cancer is classified into two major histological subtypes, including small-cell (SC) and non-small cell lung cancer (NSCLC) with NSCLC accounting for approximately 85% of cases. NSCLC is further delineated into Adenocarcinoma, Squamous cell carcinoma, and Large cell carcinoma subtypes^2^. 5-year survival rates for localized NSCLC approach 63%, but with distant metastasis the 5 year survival rates drop to 7% (American Cancer Society). Prognosis is complicated by a number of factors, including *EGFR* mutation status^3^.

A member of the ERBB family, EGFR plays a role in numerous cancers and functions through pathways PI3K/AKT, Stat3, and Ras/Raf/Mek/Erk to increase cellular growth, proliferation, and evasion of apoptotic signals. Ligand binding stimulates EGFR dimerization through conformational shifts mediated by the extracellular domains^4, 5^, resulting in a switch to the active structure. Once in the active conformation, phosphorylation occurs on the numerous tyrosine residues in the carboxy-terminal tail of EGFR^6–8^. Interestingly, specific ligands are capable of inducing differential tyrosine phosphorylation and activation of various downstream signaling pathways^9, 10^. Genetic mutations are also capable of inducing the EGFR active state, and these mutations are common in multiple cancers. Common mutations leading to constitutively active EGFR in NSCLC include a deletion in exon 19, and the L858R point mutation^11, 12^. EGFR stimulation leads to transcription of numerous gene products, from immediate early genes such as the transcription factors FOS and JUN, to secondary late response genes^13^. After signaling, EGFR is internalized and returned to the cell surface or marked for degradation^14, 15^. Interestingly, a body of literature also supports a role for EGFR in the nucleus. Indeed, EGFR has been found to act as a transcriptional activator via direct binding to A/T-rich sequences (ATRS) in the promoters of certain genes, such as cyclin D1^16^ and can act as a co-activator through interactions with transcription factors such as STAT3 to recruit nuclear EGFR to the *iNOS* promoter^17^. As a result, nuclear EGFR has prognostic value for a variety of cancers, including breast and non-small cell lung cancer^18, 19^. Taken together, EGFR is extensively involved in cancer progression through a variety of mechanisms.

With the demonstrated importance of EGFR, it is not surprising that approximately 15% of NSCLC patients have tumors presenting with amplification or activating mutations in *EGFR*, with higher percentages in Asian patients^20^. 80% of these *EGFR* mutations are putative oncogenic drivers, with the vast majority of these mutations being either missense L858R mutations or a small deletion surrounding amino acid 750, potentially resulting in an increased dimerization ability^21^. Tyrosine kinase inhibitors are standard of care for NSCLC patients who have tumors presenting with these canonical *EGFR* activating mutations. First generation Tyrosine Kinase Inhibitors (TKIs), such as Erlotinib and Gefitinib, were designed to target the ATP binding domain of EGFR. These TKIs successfully enhanced progression free survival, however resistance mechanisms develop in patients, usually in the form of a T790M *EGFR* mutation which causes a structural shift and prevents binding of TKIs to the ATP binding domain^22^. Second generation TKIs, such as Afatanib, have also been developed to target the ATP binding domain, but do so in an irreversible covalent manner. However, these second generation TKIs still suffer from resistance mechanisms due to the T790M mutation. Third generation TKIs, such as osimertinib, circumvent this structural inhibition by binding to a nearby cysteine residue and have begun to see use as first line treatment as it increases survival rates^23^. Currently, 4^th^ generation TKIs are being developed based on allosteric inhibition of EGFR to alleviate mutations associated with Osimertinib resistance. Taken together, while oncogenic mutations in EGFR are impactful, patients with these mutations have better 5-year survival outcomes due to a series of targeted tyrosine kinase inhibitors.

A critical component of EGFR activity is regulation of phosphorylation by phosphatases. A recent global screen for EGFR phosphatases revealed Protein Tyrosine Phosphatase Receptor Type H (PTPRH) as an EGFR phosphatase^24^. PTPRH, also known as Stomach Cancer-Associated Phosphatase 1 (SAP-1) is a member of the receptor like protein phosphatases. PTPRH has an extracellular region composed of several fibronectin domains, a transmembrane domain, and an intracellular phosphatase domain. The structure of PTPRH is largely conserved between humans and mice, with humans having eight fibronectin domains and mice having six^25^. In the phosphatase screening study, Yao et. al. found that PTPRH dephosphorylated EGFR, suggesting a specificity for tyrosine residue 1197.

While some phosphatases, such as PTEN^26, 27^, have well defined tumor suppressive capabilities, many phosphatases are undefined roles in the context of cancer. PTPRH studies have been largely carried out in hepatocellular tumors. Within cancers of the liver, lower PTPRH expression is associated with poorly differentiated hepatocellular carcinomas (HCC) relative to higher levels in normal liver tissue. Furthermore, overexpression of PTPRH in HCC cell lines with low PTPRH expression drastically reduced cellular motility and growth rate *in vitro*, suggesting PTPRH has a tumor suppressive role within hepatocellular carcinoma. Overexpression of PTPRH has been noted in NSCLC, with correlative hypomethylation of PTPRH being suggested as the cause^28^.

Here we have examined mutations that inactivate PTPRH, resulting in aberrant phosphorylation of EGFR using a combination of cell lines and mouse models. The role of the mutant PTPRH in NSCLC was not previously appreciated but this work illustrates that specific mutations in PTPRH may be clinically actionable using EGFR TKIs.

## Results

In our prior work involving whole genome sequencing of MMTV-PyMT FVB mice, we uncovered a conserved V483M mutation in *Pptrh* that was associated with increased phosphorylation of EGFR^29^. Examination of exome sequence from MMTV-PyMT in other background strains revealed a variety of other *Ptprh* mutations. Here we have sequenced a total of 67 PyMT mouse tumors to show the conserved V483M mutation occurs in 82% of tumors (Figure 1A). Analysis of publicly available data on other PyMT strains^30^ revealed these mutations were conserved between primary tumors and matched pulmonary metastases, suggesting that the *Ptprh* mutation occurs early in tumor progression (Figure 1B). To directly test when mutations arose in tumor progression, we extracted DNA from 21 and 35 day old MMTV-PyMT mammary glands (Figure 1C) and tested the PCR amplified sequence for presence of *Ptprh* mutations. This revealed that 9 of the 13 day 21 mammary glands and 7 of 8 day 35 mammary glands had already accumulated *Ptprh* mutations. Importantly, these samples were only hyperplastic as tumors form on average at day 45 in this background. These data suggest that there is a strong selective pressure for *Ptprh* mutations prior to overt tumor formation.

**Figure 1:**
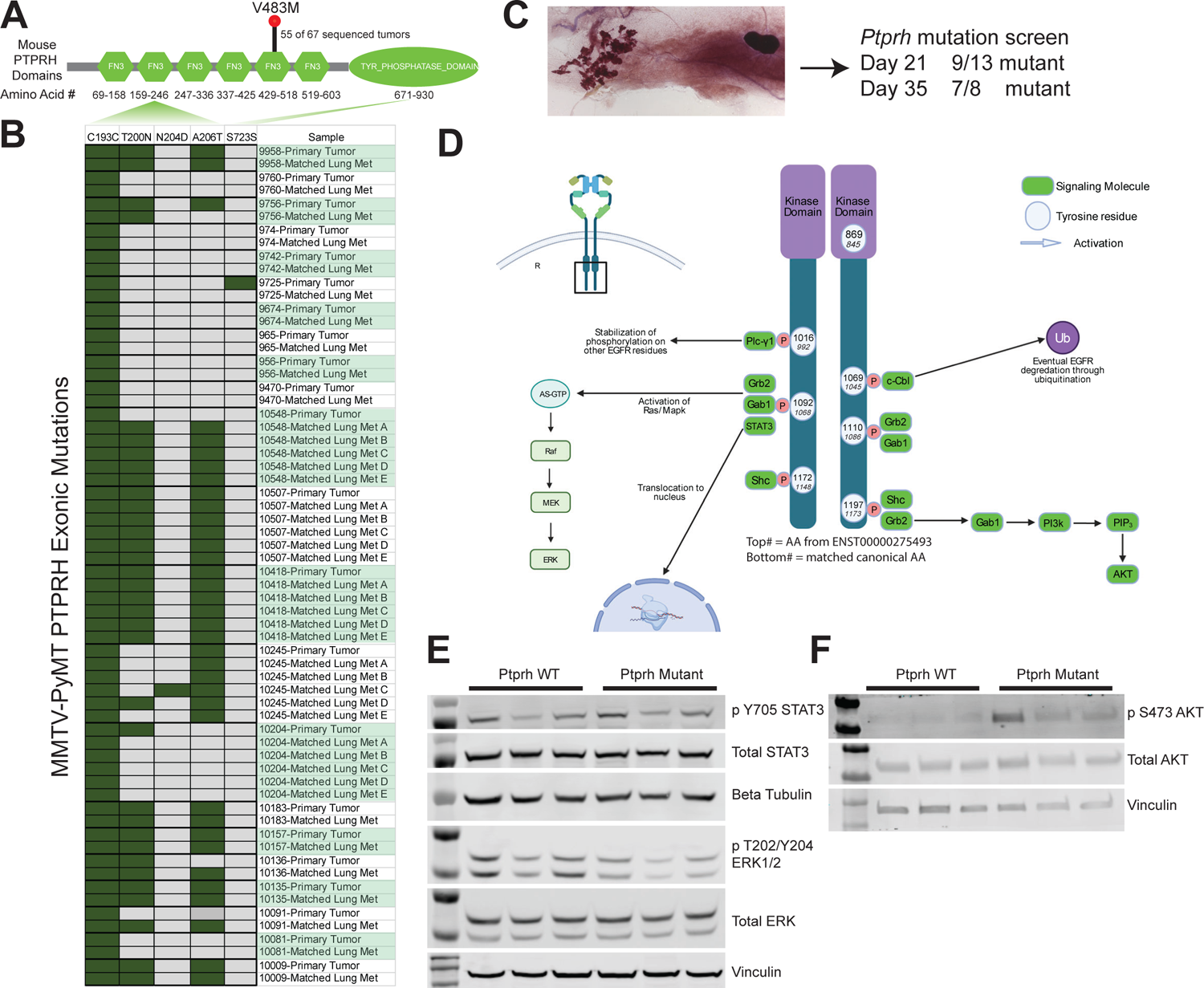
*Ptprh* Mutant Mouse Tumors have Increased Phosphorylation of AKT. Conserved metastasis and downstream regulation of EGFR pathways is seen in PyMT tumors with *Ptprh* mutations. A) Protein domain map of mouse PTPRH shows the location of conserved V483M mutated *Ptprh* within out PyMT FVB mice. B) Exome sequencing data of PyMT FVB mice from Kent Hunter lab shows *Ptprh* mutations are not conserved to one location. Furthermore, *Ptprh* mutation status is conserved between primary tumors and their matched metastasis. C) Wholemount of a day 21 MMTV-PyMT mammary gland with the hyperplastic growth on the left and the lymph node embedded in the fat pad on the right. The entire gland was used for DNA extraction and sequencing of PTPRH. D) Diagram shows the main tyrosine residues capable of being phosphorylated on the c-terminal tail of EGFR. While the diagram is not comprehensive, as signaling pathways are convoluted and undergo numerous feedback mechanisms, some of the main downstream pathways that have been characterized are shown. E) Western blotting of PyMT tumor lysates shows no increased phosphorylation of STAT3 or ERK in *Ptprh* mutant tumors as compared to WT tumors. F) Western blotting shows increased phosphorylation of AKT within *Ptprh* mutant tumors as compared to WT tumors.

Given that specificity of EGFR signaling is mediated by specific tyrosine residues (Figure 1C), we postulated that specific pathways would be activated based on which tyrosine site mutant PTPRH was failing to dephosphorylate. In Figure 1C both the canonical EGFR tyrosine residue number is listed as well as the number after the 24 amino acid signaling peptide is cleaved, as there is some confusion in the literature and available antibodies. To investigate activation of downstream pathways, *Ptprh* wild type and mutant samples were assayed for STAT3, ERK and AKT activity. No alteration to STAT3 or ERK phosphorylation was noted with *Ptprh* mutation (Figure 1D). However, mutation of *Ptprh* was associated with increased phosphorylation of AKT (Figure 1E). These data as well as our prior work with a Y1197-EGFR antibody suggest a hypothesis that PTPRH dephosphorylates Y1197 on EGFR and that *Ptprh* mutation results in an inability to downregulate signaling, leading to an increase in the PI3K / AKT signaling axis in these tumors.

To determine which human tumors contained *PTPRH* mutations, a pan-cancer search of the International Cancer Genome Consortium (ICGC) and The Cancer Genome Atlas (TCGA) data was completed (Figure 2A). Both datasets contained *PTPRH* mutations within several cancers, including a mutation prevalence of approximately 5% in non-small cell lung cancer (NSCLC). Given the incidence of *EGFR* mutations in NSCLC, and our data suggesting that mutations in mouse *Ptprh* resulted in increased EGFR activity, we hypothesized that *PTPRH* mutant human tumors would have increased EGFR signaling. To test this, we predicted EGFR activity in *PTPRH* mutant human tumors through a gene set enrichment analysis (GSEA) approach (Figure 2B). As shown in the lollipop plot, there are numerous *PTPRH* mutant NSCLC tumors with increased predicted EGFR pathway activity. These mutations were clustered within the fibronectin and phosphatase domains with localized hotspots of EGFR activity. Examining the relationship between *EGFR* and *PTPRH* mutation status from a collection of databases revealed that mutations in these two genes were mutually exclusive (p<0.0001, n=307, Figure 2C). To examine the pathways that were activated in these tumors we used single sample Geneset Enrichment Analysis (ssGSEA) on *EGFR* mutant tumors (L858R), PTPRH mutant tumors with high predicted EGFR status, and tumors that were wild type for both *EGFR* and *PTPRH* mutations. Unsupervised clustering of the ssGSEA results revealed that a subset of tumors with *PTPRH* mutations clustered together with the *EGFR* mutant tumors, suggesting a similar pathway activity profile (Figure 2D). Identity of each pathway is listed (Supplemental Table 1). Importantly, we confirmed the identification of the PI3K / AKT pathway and used GSEA to compare *PTPRH* mutant tumors to wild type, revealing a significant enrichment of the PI3K /AKT signaling axis (Figure 2E).

**Figure 2:**
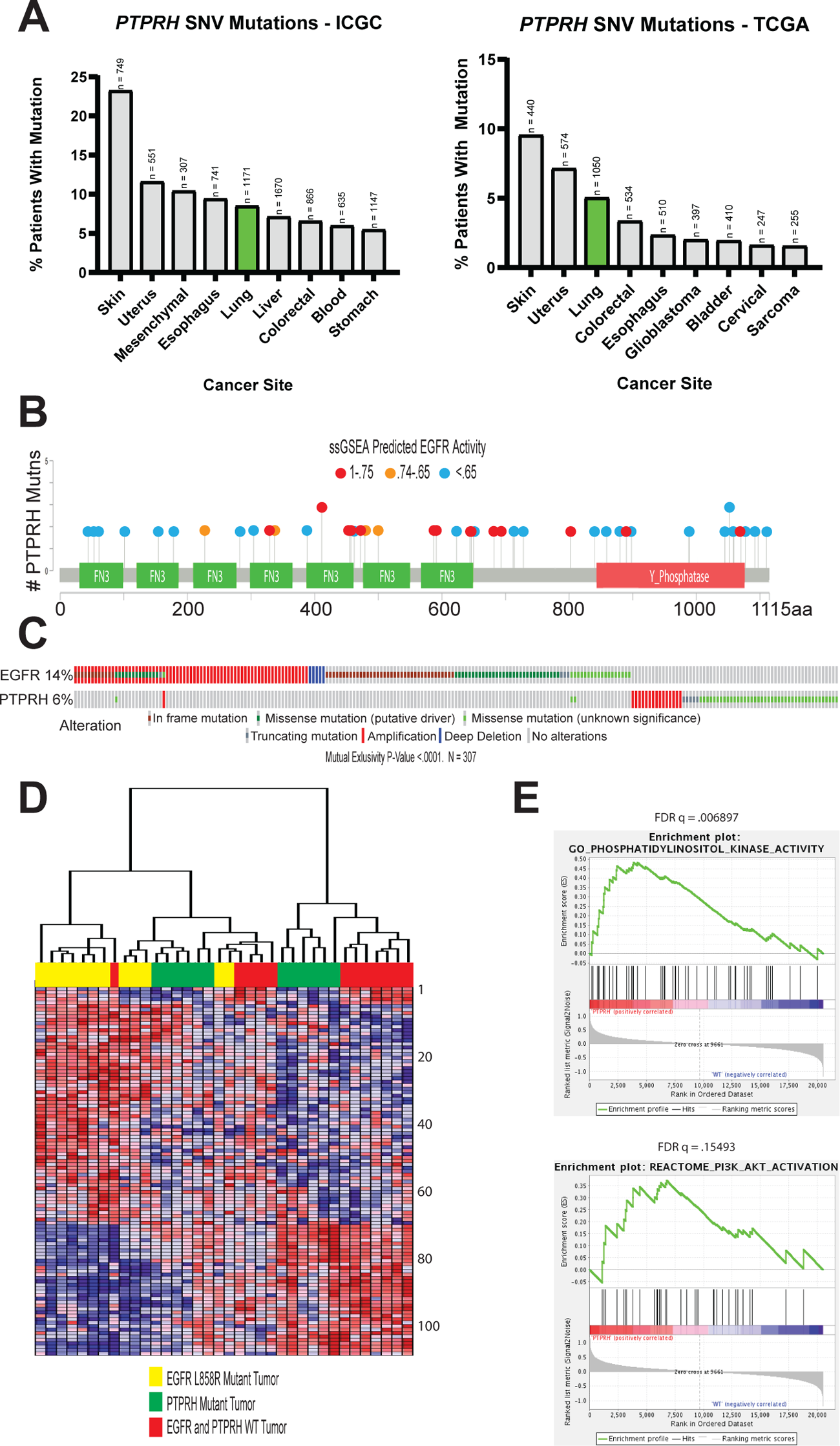
GSEA Predicts High EGFR Activity in *PTPRH* Mutant NSCLC Tumors. Numerous bioinformatics methods illustrate the importance of *PTPRH* mutations in human non-small cell lung cancer. A) data analyzed from the International Genome Consortium as well as The Cancer Genome Atlas show *PTPRH* mutations occurring in a number of human cancers. Lung cancer is highlighted due to the relationship of PTPRH with EGFR, and EGFRs importance in lung cancer. B) Lollipop plot of human *PTPRH* mutations correlated with predicted EGFR activity. Each dot represents a human NSCLC tumor with a mutation in *PTPRH*. Dot color corresponds to EGFR activity predicted through ssGSEA. C) CBIO oncoplot of NSCLC tumor mutation data from TCGA. Patient tumors with *PTPRH* mutations are shown to be mutually exclusive from patient tumors with *EGFR* mutations. D) Clustered heatmap of pathway activation prediction through GSEA. Each column represents a NSCLC tumor with mutation status corresponding to the color coded top bar. Each row represents predicted activation of pathways through ssGSEA. E) GSEA random walk plots show predicted activation of PI3K and AKT within *PTPRH* mutant tumors compared to *PTPRH* WT tumors.

Given these results were correlative in nature, we sought to directly test whether loss of PTPRH activity resulted in increased pY1197 EGFR. To test this, we used CRISPR to create knockouts of PTPRH in the H23 NSCLC cell line and flow sorted into individual clones. The generation of a knockout through insertion of an adenosine base pair at the cut site, leading to a frameshift and early stop, is shown for clone 2 (Figure 3A). Given a lack of functional antibodies for PTPRH, we instead sequenced *PTPRH* in the clonal cell lines. Importantly, we noted that several clones with an engineered knockout of PTPRH resulted in elevated levels of pY1197 EGFR (Figure 3B). To ensure specificity, we then rescued the knockout by transfecting a plasmid expressing wild type PTPRH. This restored endogenous levels of EGFR phosphorylation at Y1197 (Figure 3C). Moreover, rescue with a catalytically dead version of PTPRH (D986A) failed to result in decreased phosphorylation of EGFR (Figure 3D).

**Figure 3:**
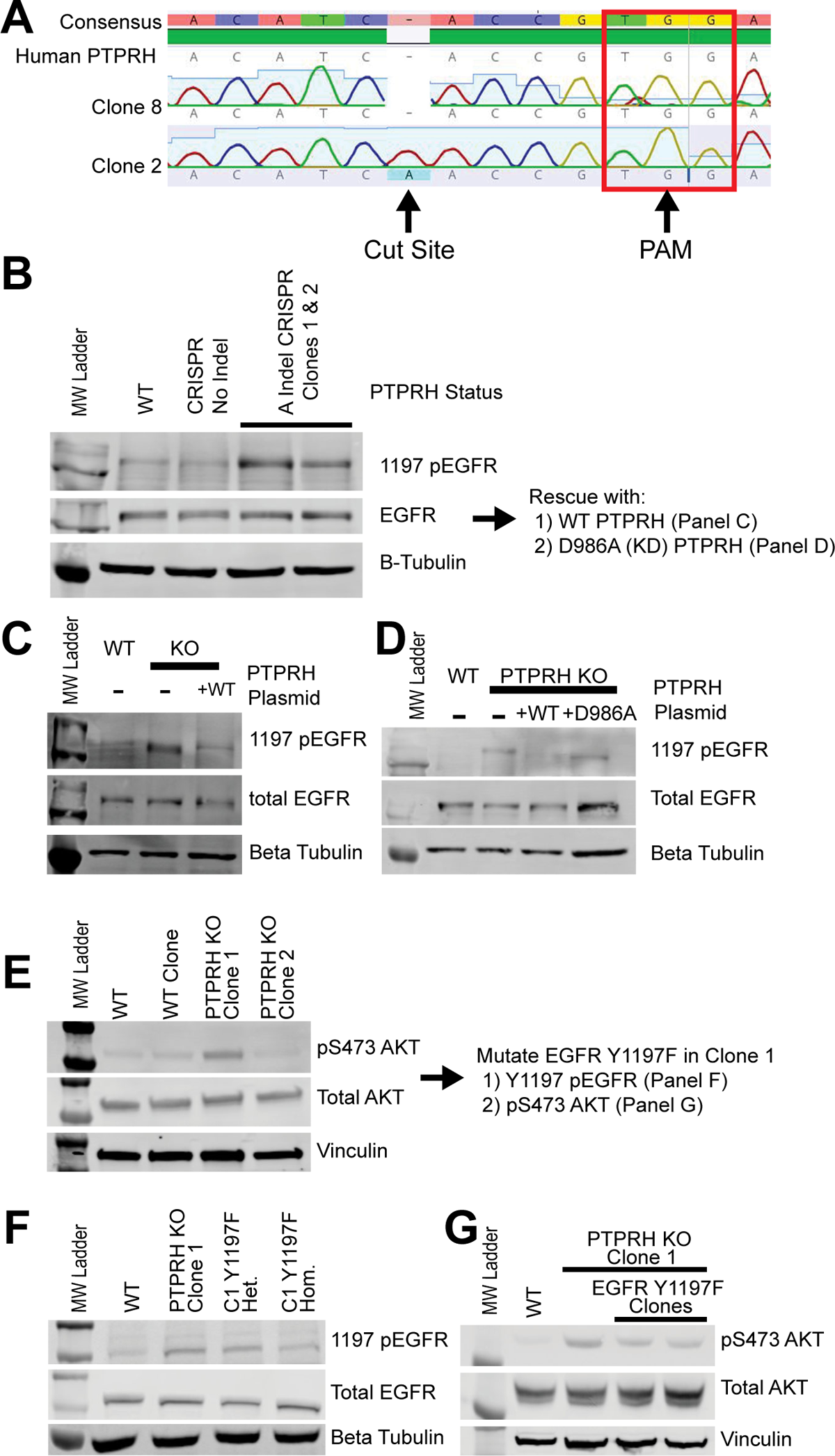
Increased 1197 p-EGFR in H23 PTPRH CRISPR KO Cells. Western blotting using lysate from PTPRH KO cells and PTPRH KO cells with transient overexpression of PTPRH plasmids demonstrates PTPRH indeed targets EGFR within human lung cancer cells. A) Electropherogram of PTPRH KO clones shows an A insertion at the CRISPR cut site. This indel was present for both clones in 3B. B) PTPRH KO CRISPR clones have increased 1197 phosphorylated EGFR. C) Overexpression of a wild type PTPRH plasmid within PTPRH KO clone 1 reduced 1197 p-EGFR. D) Overexpression of a D986A mutant PTPRH plasmid within PTPRH KO clone 1 resulted in no reduction of 1197 p-EGFR. E) Increased p-AKT is seen within 1 of the PTPRH KO clones, but not the other. F) To investigate potential clonal effects further, we created a Y1197F EGFR mutation within H23 PTPRH KO clone 1. G) Step-wise reduction of 1197 EGFR phosphorylation is seen within heterozygous and homozygous Y1197F clones. These Y1197F clones also have marked reduction in p-AKT.

Given that both the mouse and human computational predictions suggested a role for the PI3K / AKT pathway but not the ERK or STAT3 pathways, we examined the various PTPRH knockout lines for phosphorylation of these downstream pathways. As expected, no alterations were noted in the ERK or STAT3 pathways (Supplemental Figure 1). In contrast, AKT phosphorylation was noted but was variable between the knockout clones (Figure 3E). Given the potential for clonal effects, Y1197F mutations in EGFR were engineered in the PTPRH KO clone with elevated pAKT. Both heterozygous and homozygous Y1197F EGFR mutations were examined and a step-wise reduction in EGFR Y1197 phosphorylation was detected (Figure 3F). Within the two Y1197F clones we also noted a reduction in pAKT, consistent with the hypothesis that Y1197 was the target of PTPRH (Figure 3G).

Due to the potential for clonal effects that would impact analysis, pooled knockouts of *PTPRH* were generated in the H23 cell line. Tracking of Indels by Decomposition (TIDE)^31^ analysis revealed a knockout efficacy of 45% (Supplemental Figure 2). As expected, Western blotting revealed increased phosphorylation of EGFR at Y1197 in this pooled line (Figure 4A). Loss of PTPRH was also associated with increased growth rate and proliferation in growth curves (Figure 4B) and MTT assays respectively (Figure 4C).

**Figure 4:**
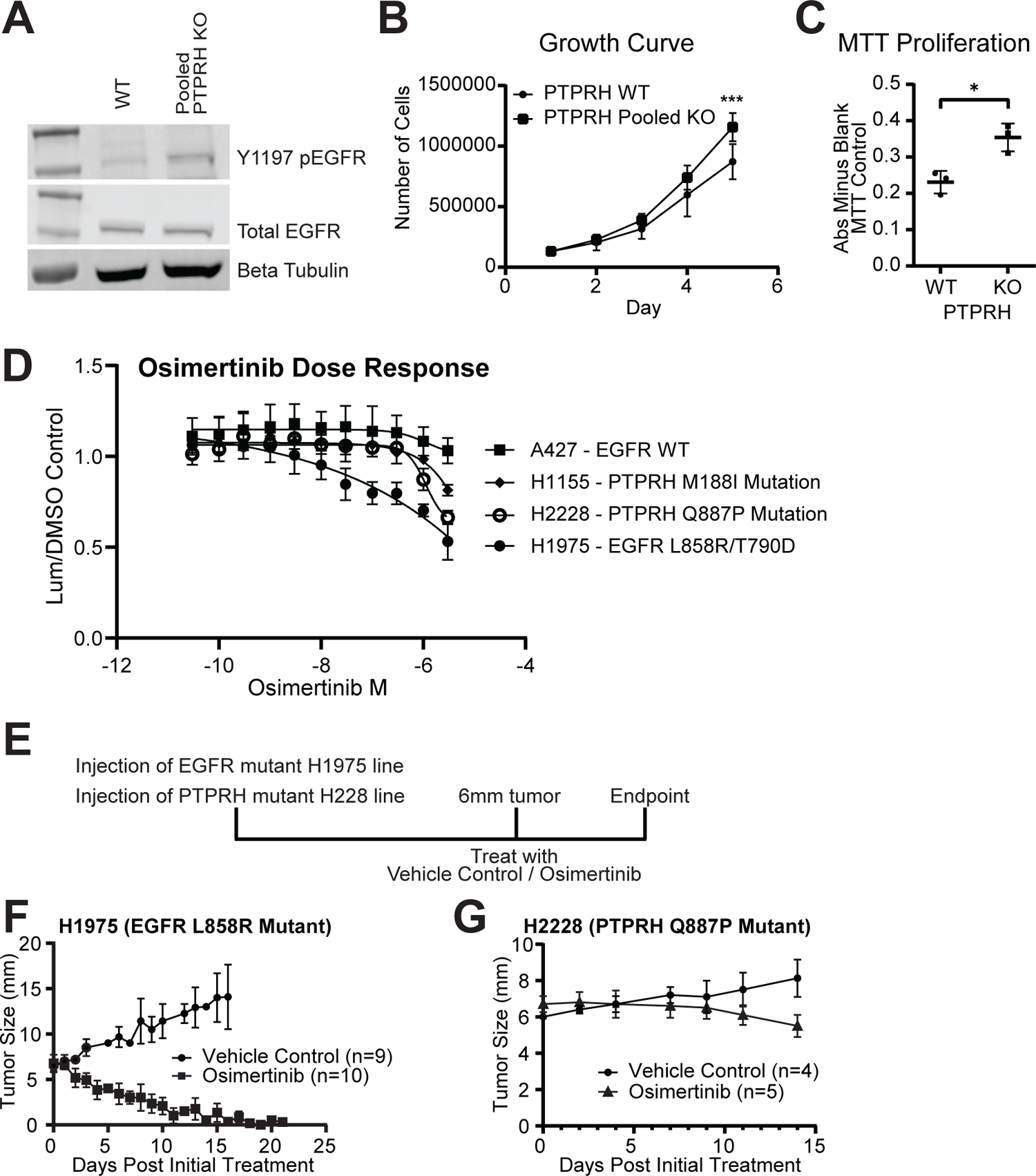
*PTPRH* mutant cell lines respond to TKI osimertinib Pooled PTPRH KO cells have increased proliferation, and *PTPRH* mutant cell lines respond to osimertinib. A) Western blotting confirms increased p-EGFR at tyrosine 1197 within pooled KO cells compared to WT cells. B) Cellular growth curves show increased growth in PTPRH pooled KO cells as compared to WT cells. C) MTT assays completed with H23 WT and H23 PTPRH pooled KO cells show increased proliferation. D) Two *PTPRH* mutant cell lines (WT for EGFR) from human non-small cell lung cancer tumors show response to the TKI osimertinib *in vitro*. E) Treatment plan for *in vivo* treatment of H2228 *PTPRH* mutant cell line. Either H1975 (L858R EGFR mutant) or H2228 (Q887P *PTPRH* mutant) cells were injected into the left flank of nude mice. Mice were then randomized into two treatment groups, vehicle control or osimertinib. H1975 mice were treated with 25 mg/kg of osimertinib and H2228 injected mice were treated with either 25 mg/kg or 50 mg/kg of osimertinib. F) *in vivo* drug curve showing response to osimertinib of H1975 *EGFR* mutant injected mice. G) *in vivo* drug curve showing response to osimertinib of H2228 *PTPRH* mutant injected mice.

Given that loss of PTPRH resulted in elevated phosphorylation of EGFR, we hypothesized that these tumors would be susceptible to EGFR TKIs, demonstrating this in a proof of principle experiment in PyMT tumors^29^. To test this hypothesis in human lung cancer cell lines, dose response curves were completed for four cell lines using the EGFR TKI Osimertinib. NSCLC lines A427 (WT for EGFR and PTPRH) was a negative control while the H1975 line with classic L858R / T790D activating EGFR mutations was a positive control. Two NSCLC lines with naturally occurring *PTPRH* mutations were also tested including H1155 with a M188I mutation in the second fibronectin domain and H2228 with a Q887P mutation in the phosphatase domain. Both of these spontaneous mutations were predicted to have moderate EGFR activation and were not in the regions with the highest EGFR predicted activity for *PTPRH* mutations. As expected, the A427 negative control had no response while the H1975 positive control had a robust response. Interestingly, the two PTPRH mutant NSCLC lines had an intermediate response (Figure 4D). To test whether this would have phenotypic effects *in vivo*, we used a strategy of injecting H2228 cells into the flank of mice and treating with the EGFR TKI osimertinib once tumors reached 6mm (Figure 4E). The positive control responded well with tumors rapidly shrinking with TKI treatment (Figure 4F). While slower growing, the H2228 line with the *PTPRH* mutation had an appreciable tumor reduction at 50 mg/kg dosage (Figure 4G) but had no effect at 25 mg/kg (data not shown). The study was stopped at 14 days due to endpoint concerns. While not as robust as the EGFR mutant result, these proof of principle data demonstrate that *PTPRH* mutation status can induce a susceptibility for EGFR TKIs.

To determine how widespread EGFR activation was within *Ptprh* mutant mouse tumors, and *PTPRH* knockout human tumors injected into the flank of mice, we employed an immunohistochemistry approach using an antibody specifically recognizing phosphorylation at Y1197 in EGFR. In a negative control (MMTV-PyMT) that was wild type for *Ptprh* mutations, we noted no appreciable staining for pY1197 EGFR (Figure 5A and B). In MMTV-PyMT tumors with a *Ptprh* mutation, there was widespread staining for pY1197 EGFR (Figure 5C). Interestingly, higher magnification revealed that this staining was predominantly nuclear (Figure 5D). Examining human NSCLC in the H23 PTPRH knockout line revealed increased pY1197 EGFR staining as compared to wild type controls (Figure 5E), but this staining was noted to be primarily located on the membrane (Figure 5F).

**Figure 5:**
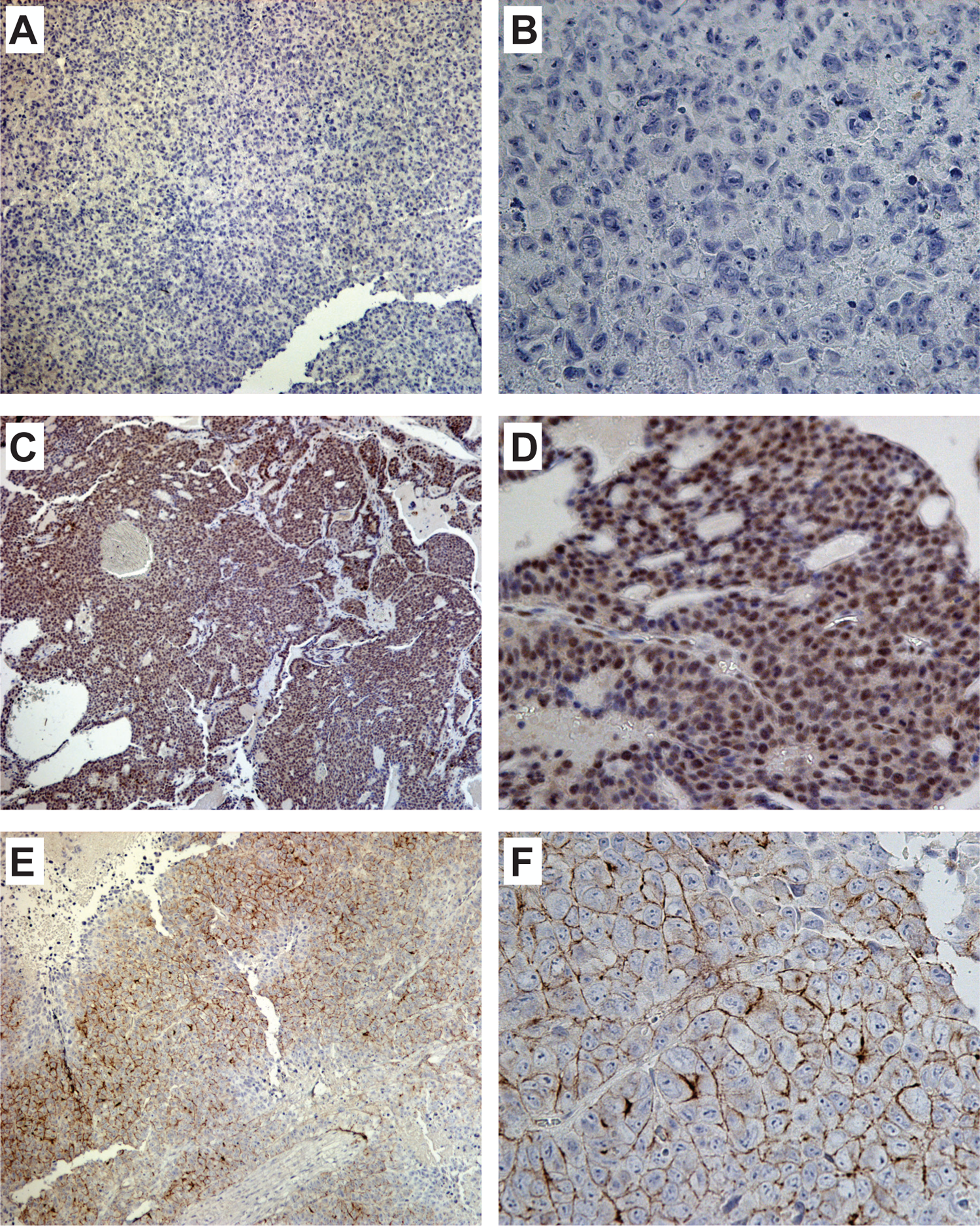
Phospho-EGFR Immunohistochemistry reveals nuclear and membrane staining. An MMTV-PyMT tumor that was wild type for Ptprh was used in immunohistochemistry for phosphoEGFR revealing essentially no staining at low (10x) and high (40x) magnification (A and B respectively). A PyMT tumor with a V483M PTPRH mutation revealed largely nuclear staining across the entire tumor and was reflective of these tumors. Staining the H23 PTPRH knockout human tumor line grown in mice revealed membrane specific staining for phosphoEGFR (E and F).

The activity of PTPRH is likely not limited to Y1197 of EGFR. To determine what other kinases were regulated by PTPRH, we screened a phosphorylated receptor tyrosine kinase (RTK) array with lysate from wild type H23 cells (Figure 6A) and lysate from CRISPR PTPRH KO H23 cells (Figure 6B). Several RTKs with differential phosphorylation patterns with PTPRH knockout lysate were identified, including FGFR1 with a 3.8 fold increase and IGF-1R with a 2.4 fold increase (Figure 6A-C). Examining these RTKs in the publicly available genomic data through FGFR1 and IGF-1R ssGSEA signatures revealed predicted activation of these kinases in many of the mutations that also resulted in EGFR activation, although others were unique to each RTK (Figure 6D). To confirm predicted activation of FGFR1, Westerns were completed using cell lysates from PTPRH WT and PTPRH KO H23 cells revealing a clear increase in phosphorylation of FGFR1 within PTPRH KO cells (Figure 6E).

**Figure 6:**
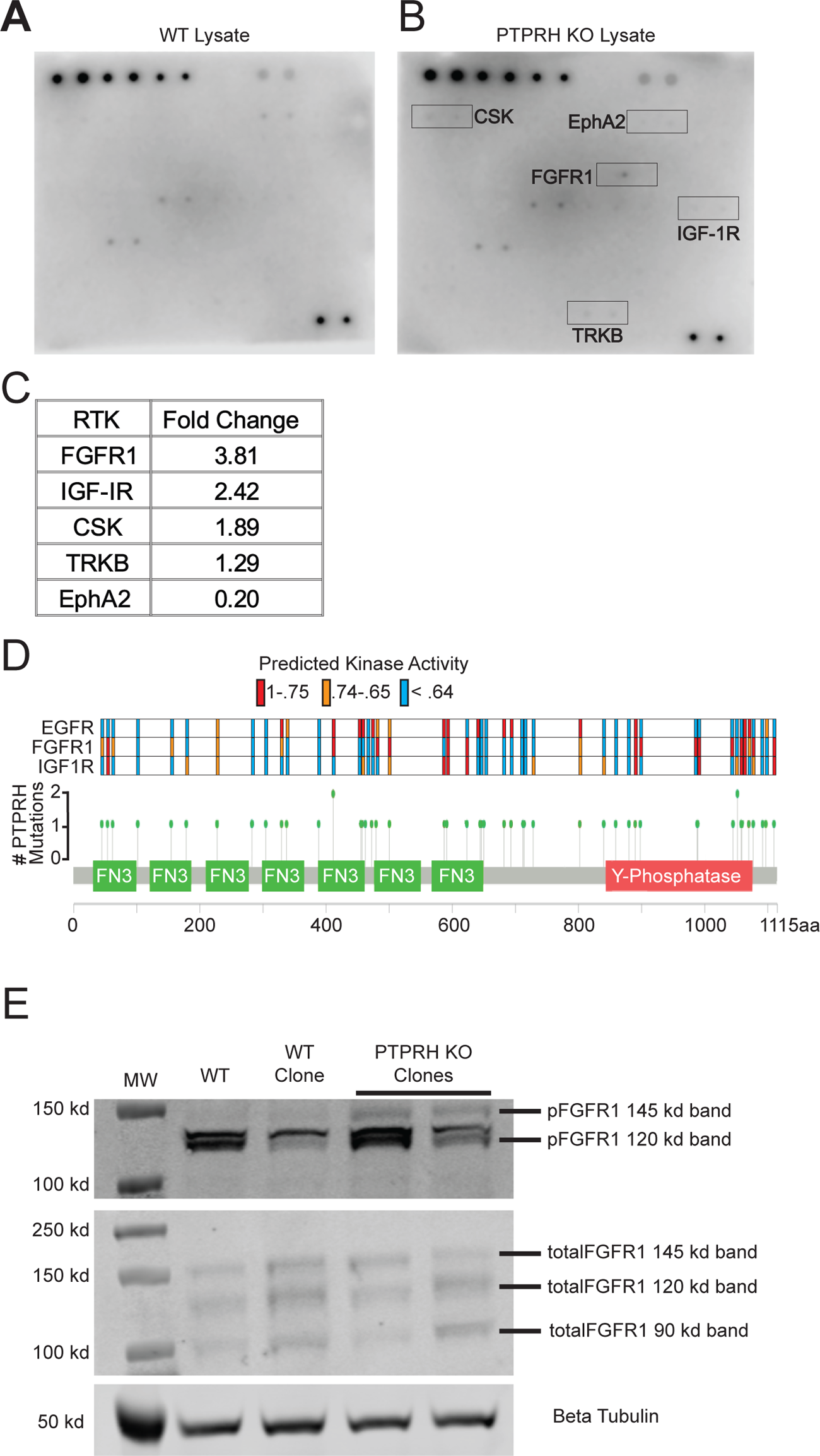
PTPRH Regulation of Receptor Tyrosine Kinases outside the ERBB Family Ablation of PTPRH results in differential activation of numerous receptor tyrosine kinases outside of EGFR. A) A human phosphorylated RTK array shows differential phosphorylation of numerous RTKs when incubated with lysate from H23 PTPRH WT cells. B) A human phosphorylated RTK array shows differential phosphorylation of numerous RTKs when incubated with lysate from H23 PTPRH KO cells. Top five differentially phosphorylated RTKs are highlighted in red. C) Table showing the RTKs with the top 5 largest fold changes between PTPRH KO and PTPRH WT lysates. D) The lollipop plot in figure 2B was recreated, adding predicted activation for FGFR1 and IGF-1R. Briefly, each dot on the PTPRH exome plot corresponds to a *PTPRH* mutant NSCLC tumor. Color-coded bars above each dot correspond to the predicted activity of EGFR, FGFR1, or IGF-1R. E) Western blotting confirms increased phosphorylation of FGFR1 at tyrosine residues 653/654, within PTPRH KO cell lysates compared to PTPRH WT lysates.

## Discussion

Here we have identified conserved V483M *Ptprh* mutations in mouse mammary tumors from MMTV-PyMT transgenic mice. In a mixed background these mutations were conserved in matched pulmonary metastases, indicating this mutation occurs early in tumorigenesis. We demonstrated mutation of *Ptprh* to be impactful since PTPRH no longer dephosphorylated Y1197 of EGFR, resulting in activation of the PI3K / AKT signaling cascade. Given the importance of EGFR activity in lung cancer, we confirmed the causal nature of PTPRH loss on EGFR activity through CRISPR mediated knockout of *PPTPRH* in a NSCLC line by observing increased pEGFR and pAKT. Importantly, a rescue experiment demonstrated that this was a specific event as plasmid expressed PTPRH was able to dephosphorylate EGFR while the rescue with a catalytically dead PTPRH did not. In addition, loss of PTPRH resulted in increased growth rate, potentially as a function of activation of the EGFR / PI3K / AKT signaling pathway. These results are summarized in Figure 7. This has potential to be a tumor driving event as it occurs early in tumor etiology and allows activation of a major signaling pathway to inappropriately persist. The conservation of the V483M mutation in over 80% of tumors from the genetically engineered mice in the FVB background also indicates that there is a remarkable selective pressure for EGFR pathway activity. A pan-cancer analysis of human *PTPRH* mutations found numerous cancers harboring mutations, suggesting mutated PTPRH may play a role in tumor development across the spectrum of cancer types. *PTPRH* mutations were found in approximately 5% of NSCLC patients, with these mutations spread across the PTPRH exome. This is an interesting contrast to the conserved V645M mutation found within PyMT tumors, with potential implications for which mutations may be impactful on tumor growth. While a mechanism has yet to be explored for each of the various mutations, it is likely that the mutations are acting in a different fashion from each other. Mutations within the phosphatase domain may abrogate catalytic activity, while mutations in the fibronectin domains may prevent dimerization and binding of target substrates^25^. Since some of the phosphatase domain mutations with predicted high EGFR activity lie outside the conserved phosphatase activity HC(X_5_)R motif, it is also possible that these mutations impact access to the phosphatase domain and prevent recognition of substrate binding sites. Given the lack of conserved human *PTPRH* mutations, eventual utility of *PTPRH* mutation screening would need to be combined with a functional output screen. Ultimately the mutations in PTPRH and their functional impact on EGFR and response to TKIs may be paired with pEGFR or pFGFR1 status to predict response to EGFR TKI.

**Figure 7:**
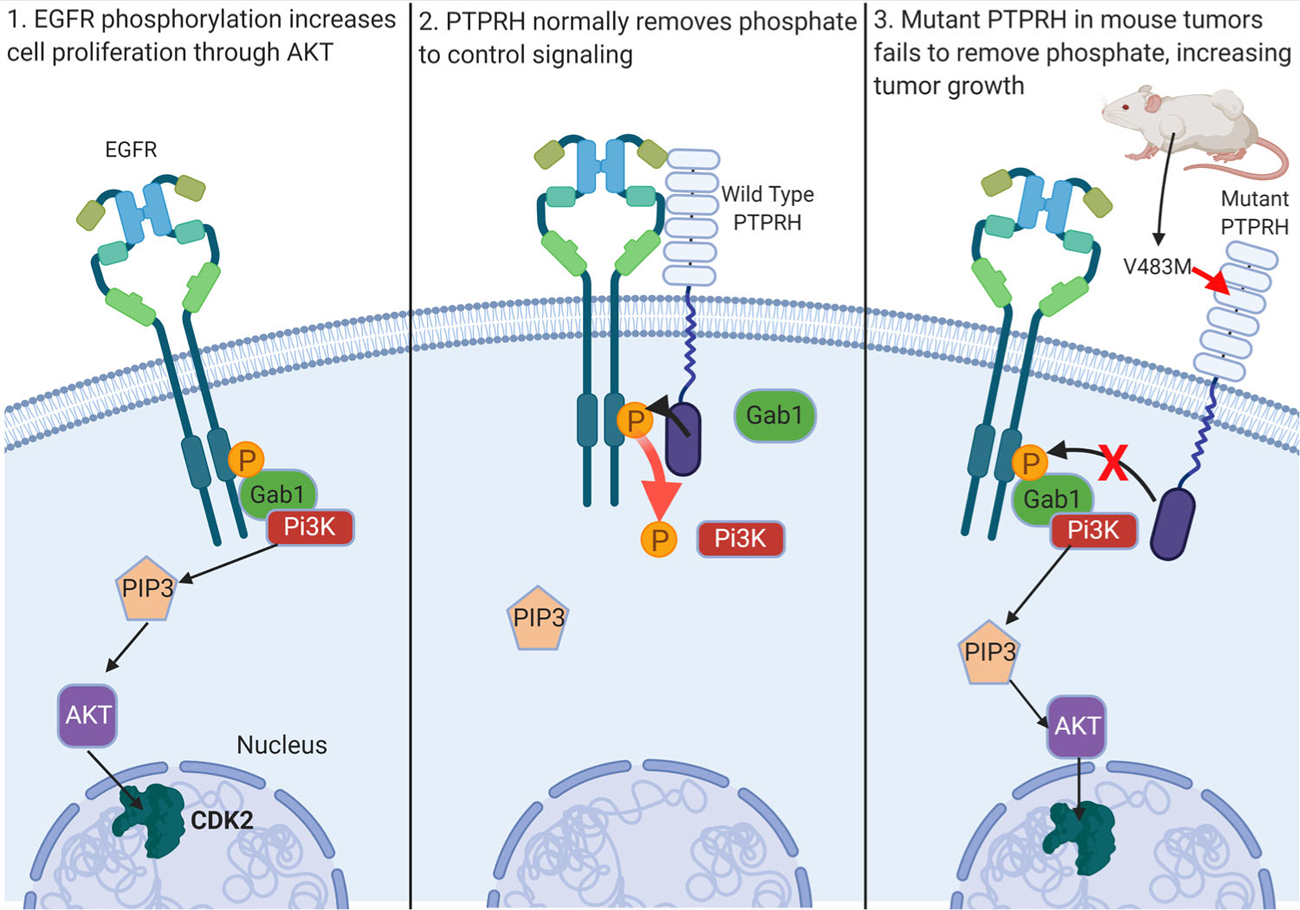
Schematic of *Ptprh* mutant mouse tumors failing to dephosphorylate EGFR. Ordinarily, PTPRH is responsible for regulating EGFR signaling through dephosphorylation of tyrosine residues on the C-terminal tail of EGFR. We have shown PTPRH mutant mouse tumors and PTPRH KO human cells to have increased phosphorylation of EGFR, and subsequent increased activation of the PI3K/AKT pathway.

Our findings demonstrated increased phosphorylation of EGFR upon loss of PTPRH in NSCLC in both *in vitro* and *in vivo* models. With this causality and given that 5% of NSCLC tumors have mutations in *PTPRH*, and an estimated 235,000 cases of lung cancer occurring yearly within the United States, over 10,000 patients who present with *PTPRH* mutations could potentially benefit from EGFR targeted TKI therapy. Interestingly, the mutation types may allow for prediction of the therapeutic response. For instance, here we identified two NSCLC lines with *PTPRH* mutations that only had a medium level of predicted EGFR activity and treated them with the TKI Osimertinib. While both responded *in vitro*, the H2228 cell line also responded *in vivo*, with a decrease in tumor volume. In the future, cataloging PTPRH mutations with EGFR TKI response would allow for appropriate clinical action. Other potential options for treatment of PTPRH targets include dual inhibition of kinases whose signaling pathways are altered by PTPRH loss, or targeting RTKs with proteolysis targeting chimera (PROTAC) molecules, which target them for degradation. Overall, treatment of downstream targets regulated by phosphatases, rather than the phosphatases themselves, may be a viable solution, although this will would require considerable characterization of the pathways affected by deregulated phosphatases. This is especially important to consider with the context dependent nature of PTP regulation, such as PTPRH deactivating EGFR.

Beyond EGFR, a kinase array showed increased phosphorylation of numerous RTKs within PTPRH knockout cell lines, including FGFR1 and IGFR1. Interestingly, increased phosphorylation of EGFR was not detected on the array. However, a closer examination of the phosphorylated antibodies used in the array revealed that EGFR Y1197 was not included. The increased phosphorylation of FGFR1 was further confirmed through western blotting. The activation of the FGFR1 pathway has interesting implications, both at the level of cellular pathways that may be affected, as well as potential treatment options for those with non-functional PTPRH. Moreover, the role of FGFR1 in other cancers has potential to open EGFR therapy in the other tumor types. In addition, with further preclinical work, a dual drug inhibition approach of targeting FGFR1 and EGFR may be of clinical use.

Finally, *Ptprh* mutant mouse tumors have increased staining of nuclear EGFR. Nuclear EGFR has been noted in times of cellular stress, as well as regenerating liver tissue. While in the nucleus, EGFR can act as a cofactor, or direct transcriptional activator by binding to the promoters of certain genes, such as cyclin D1. Increased nuclear EGFR upon loss of PTPRH activity could have profound impacts on cellular signaling pathways. The mechanism behind increased nuclear localization of EGFR has not been explored but warrants further exploration.

## Methods

### Targeted Resequencing of PyMT tumors

DNA was extracted from flash frozen tumors using lysis buffer (50 mL Tris HCl, 5 mL 500 mM EDTA, 10 mL 10% SDS, 20 mL 5M NaCl, H20 up to 500 mL), or FFPE tissue using the Qiagen FFPE extraction kit. The region flanking V483M was PCR amplified using the following primers; Forward 5’ GGCCTTAGGTTCAATTGTGAATAC 3’ Reverse 5’ CCTTAGCTTCCCGAGTATTGGTT 3’ Amplified DNA was sent to GeneWiz for Sanger sequencing with the following primer 5’ TCATCCAAACTACATCTATGATCCA 3’. Geneious software (https://www.geneious.com/) was used for alignment to reference DNA.

### Analysis of PTPRH Mutations in Exome Sequence Data

Pre-annotated VCF files were downloaded for 64 tumors from GEO ascension number GSE142387. Data was processed within R by reading in VCF files, then filtering to only keep mutations within the Chr 7 bp 4548992 – 4604041 range (location of *Ptprh* in mouse genome). These files were then converted to Annovar format, exported, and annotated using Annovar in Linux based command. Statistical analysis was completed using a student’s t test (unequal variance, 2 tailed) between the metastasis group (mutations per met sample), and the primary group (mutations per primary tumor).

### PTPRH Mutations in Human Cancers

Pan-Cancer datasets from numerous sources, including TCGA and ICGC, were analyzed through CBioPortal and the ICGC portal. Lung cancer mutation percentage were analyzed specifically using TCGA 2016 dataset accessed through CBioPortal (https://www.cbioportal.org/). The South Korean and U.S datasets showing discrepancy in percentage of *PTPRH* mutations were analyzed on the ICGC portal (https://dcc.icgc.org/). Both datasets were filtered to include only patients with exonic mutations.

All NSCLC datasets available on CBioPortal were used for mutual exclusivity analysis and are listed below. *PTPRH* and *EGFR* SNV mutation data were downloaded and combined. Duplicate samples were removed, and any sample with a *PTPRH* or *EGFR* mutation was considered. A 2×2 contingency table was run to determine mutual exclusivity. Datasets include; MSK - Cancer Cell 2018, MSKCC - J Clin Oncol 2018, TRACERx - NEJM 2017, University of Turnin, 2017, MSK - Science 2015, TCGA - Nat Genet 2016 (Pan), Broad - Cell 2012, MSKCC - Science 2015, TCGA - Firehose Legacy, TCGA - Nature 2014, TCGA - Pan-cancer Atlas, TSP - Nature 2008, MSKCC - Cancer Discov 2017, TCGA - Nature 2012

### Demographics of PTPRH mutations

Age, overall survival, and race demographics were analyzed using the Lung Adenocarcinoma TCGA Pan-Cancer Atlas data set downloaded from CBioPortal. Two-tailed Student’s T-Tests assuming unequal variance were completed for *PTPRH* mutant VS. *EGFR* mutant samples, as well as *PTPRH* mutant VS. WT (non-EGFR mutant) samples for age of diagnosis and overall survival. Samples without age or OS data were excluded. Only samples with missense or truncating mutations were included, and overexpression samples were excluded. Race was analyzed using a 2×2 contingency table.

### EGFR Activity and pathway activity predictions

TCGA pan-cancer RNA-seq dataset (downloaded from UCSC Xena) was analyzed for *PTPRH*, *EGFR*, *FGFR1*, and *IGF1R* mutations. This mutation list was downloaded and filtered to keep samples that had a mutation in *PTPRH*, *EGFR*, or that were wild type for *PTPRH*, *EGFR*, *FGFR1*, and *IGF1R*. Any sample with a mutation in *PTPRH* was kept, resulting in 53 samples. 10 samples of each of the two categories were kept; WT for *PTPRH* and the above three RTKs, and L858R mutant *EGFR* that were WT for *PTPRH*, *FGFR1*, or *IGF1R*. To decide which WT and EGFR samples to keep, samples from those subsequent groups were assigned a random number using the RAND() function in Excel. These numbers were then sorted from highest to lowest, keeping the top 10 samples. RSEM(log2 X+1) normalization was applied to the filtered sample list, resulting in 47 *PTPRH* mutant samples (WT for the kinases), 9 samples that WT for *PTPRH* and the three kinases, and 8 samples with *EGFR* mutations (WT for *PTPRH*, *FGFR1*, and *IGF1R*). ssGSEA was run on the samples to predict pathway activation status. Pathways for each kinase were filtered down, selecting the most relevant and robust pathway. A ranking sum score was applied to the pathway prediction data for each sample.

For GSEA analysis of *PTPRH* mutant tumors, the pan-cancer RNA-seq dataset was again downloaded from UCSC Xena. Twelve tumors for each of the three categories were kept; *PTPRH* mutant tumors predicted to have high EGFR activity, *EGFR* L858R mutants, and tumors that were WT for both *PTPRH* and *EGFR*. GSEA was completed using the GenePattern server.

### CRISPR Knockout

Benchling was used to design the guide RNA (AGCACACACTAACATCACCG) targeting the fourth exon of *PTPRH*. The guide was cloned into px458 using AgeI and EcoRI. Transient transfection of px458 into H23 cells was completed using Promega’s Viafect. GFP positive cells were sorted into single cell clones into 96 well plates using FACS. Once clones had grown into a colony, they were subsequently moved to 24-well plates, then 6-well plates. DNA was harvested and sent to ACTG for sanger sequencing.

### CRISPR Knock-In Mutations

Guide RNA was designed in Benchling with the PAM (NGG) sequence 5 bp downstream of the desired EGFR Y1197 mutation site. The single stranded region of homology was designed in Benchling by choosing desired length for homology arms as well as the desired mutation, then taking the reverse complement of that strand. The oligo was designed with 36 bp upstream of the desired mutation site and 90 bp downstream. The desired mutation resulting in a Y1197F amino acid substitution was added. This mutation also resulted in the addition of an EcoRI cut site, which was used for downstream screening. The mutation also altered the guide RNA enough to prevent re-annealing once HR mediated repair occurred. Guide RNA was cloned into px458. H23 PTPRH KO cells were transfected using Viafect in a 6:1 ratio. 1 ug of px458 with guide, and 4 ug of ss repair template were transfected. Sorting was completed using FACS for GFP. Clones were screened using a digest for EcoRI and confirmed with sequencing.

### Western Blotting

Tumor lysates were harvested from flash frozen tumors by crushing with a mortar and pestle, then dissolving in TNE lysis buffer (5 mL 1 M Tris HCl pH 8, 3 mL 5M NaCl, 1 mL NP40, 400 uL .5M EDTA, 2.0 mL .5M NaF, H2O to 100 mL). Roche mini protease tablets and sodium orthovanadate were used as protease and phosphatase inhibitors respectively. Primary antibodies were incubated overnight. Antibodies used were as follows; total EGFR (Cell Signaling D38B1), 1197 EGFR (Invitrogen PA5-37553), total AKT (Cell Signaling 11E7), p-s473 AKT (Cell Signaling D9E), total STAT3 (Cell Signaling 79D7), p-Y705 STAT3 (Cell Signaling D3A7), total FGFR1 (Cell Signaling D8E4), p-Y653/654 FGFR1 (Cell Signaling 3471s), beta tubulin (Proteintech 10094-1), vinculin (Cell Signaling E1E9V), total ERK (Cell Signaling 9102), p-ERK (Cell Signaling 4370).

### Overexpression

PTPRH cDNA within plasmid PRc-CMV was kindly provided by Dr. Takashi Matozaki at Kobe University. Site directed mutagenesis was used to achieve a D986A mutant. Both WT and D986A mutant *PTPRH* plasmid constructs were transiently expressed in PTPRH KO cells using Viafect.

### RTK Array

The manufacturer’s protocol for RayBiotech Human RTK Phosphorylation Array C1 kit was followed. Membranes were incubated with lysate from H23 WT cells or H23 PTPRH KO cells.

### IHC Nuclear EGFR

Human cell lines H23 PTPRH WT or H23 PTPRH KO were injected into the left flank of nude mice. H23 cell line tumors were grown to approximately 10 mm in the largest direction prior to necropsy. Mouse PyMT tumors, and tumors grown from human H23 cells were necropsied with portions of tumor tissue preserved in formalin, and portions of tumor flash frozen for further downstream analysis. Formalin fixed paraffin embedded tumors were subjected to staining using an antibody specific for 1197 EGFR (Thermo PA5-37553).

### Pooled CRISPR Knockout

Guide RNA (AGCACACACTAACATCACCG) for PTPRH was designed using Benchling. And cloned into a lentiviral Cas9 plasmid (Addgene # 52961). Viral generation was completed through transfection of 293T cells with packaging plasmid psPAX2 and envelop plasmid pMD2.G in a ratio of 3.7:1.2:5 with the Cas9 plasmid respectively. Viral supernatant was collected from 293T cells 3 days after transfection, and filtered through a .22 uM syringe filter. 1 mL of filtered viral supernatant was applied to H23 WT cells at ∼30% confluency. Sanger sequencing was used to confirm knockout, and for TIDE analysis.

### MTT Assay and Growth Curves

MTT assay kit (Roche 11465007001) instructions were followed. Graphpad was used to plot and statistically analyze results. A Welch’s two-tailed t-test yielded a p-value of .0137.

For growth curves 1.0 x 10^5^ cells were plated in triplicate within 6-well plates. On days 1-5, cells were trypsinized and cell number was read using an automated cell counter. Graphpad was used to plot results.

### Dose response curves

Cells were diluted to 5.0 x 10^4^ cells per mL, and 20 uL of cell suspension was added to wells of an opaque 384 well plate. After overnight recovery, cells were subjected to a dose response curve of increasing drug concentration in half log steps. For single drug curves, osimertinib (Cayman AZD9291) range was .00003 to 30 uM. For dual drug curves, osimertinib range was .03 to 10 uM, and either KRAS inhibitor (ARS853, Cayman) or FGFR1 inhibitor (PD166866, Cayman) range was .00003 to 30 uM. 10 mM stocks of drugs were made by diluting with DMSO, and half-log drug series were diluted fresh with complete media. Cell viability was read after 48 hours using Promega’s Cell Titer Glo. Luminescence values were normalized to non-drug treated controls, and plotted using Graphpad.

### *In vivo* treatment

H2228 and H1975 cell lines were injected into the left flank of 6-12 week old nu/nu mice. After tumors reached 6mm in the largest dimension, mice were randomized into treatment groups; vehicle control, 25 mg/kg osimertinib, or 50 mg/kg osimertinib. The 50 mg/kg dose was only used for mice with H2228 tumors. Osimertinib (AZD9291 Cayman) was diluted using the following in order to achieve a final ratio: 5% DMSO, 40% polyethylene glycol, 5% tween-80, 50% H2O. Max volume of treatment was 10 uL for 1 gram of body weight. Mice were weighed on first day of treatment, and volume of drug was adjusted to achieve proper dose.

### KI67 scoring

Slides were scored on a scale of 1-10 by three blinded reviewers independently. The mean for each tumor slide was then taken by averaging the three reviewer scores for each slide. A two-tailed student’s T-test assuming unequal variance was then completed across the osimertinib and vehicle control sample groups, using the means for each tumor slide (each group n=4).

## Acknowledgements

We appreciate the plasmid that was a gift of Dr. Matozak.

---

**Supplemental Figure 1.**
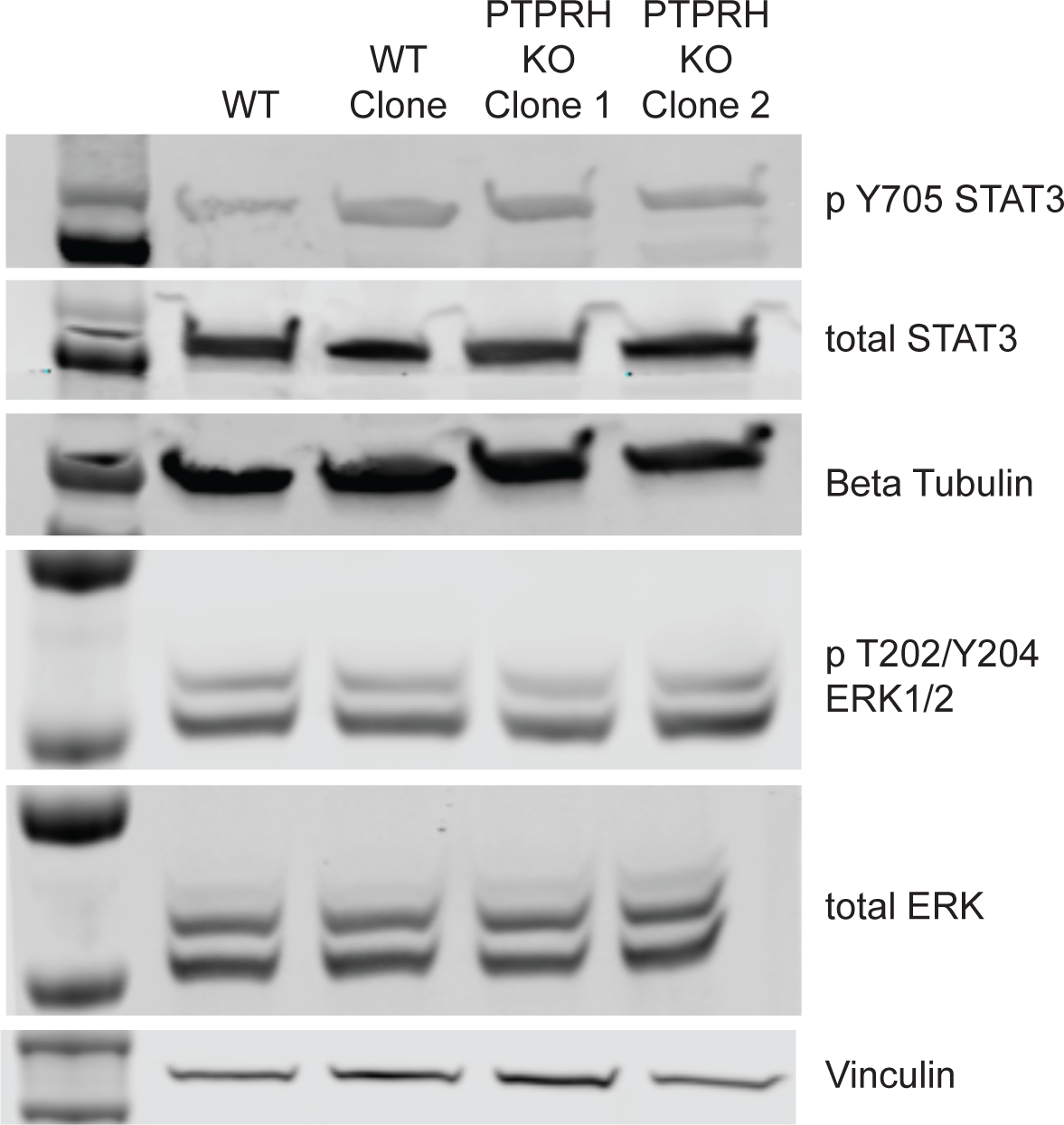

**Supplemental Figure 2.**
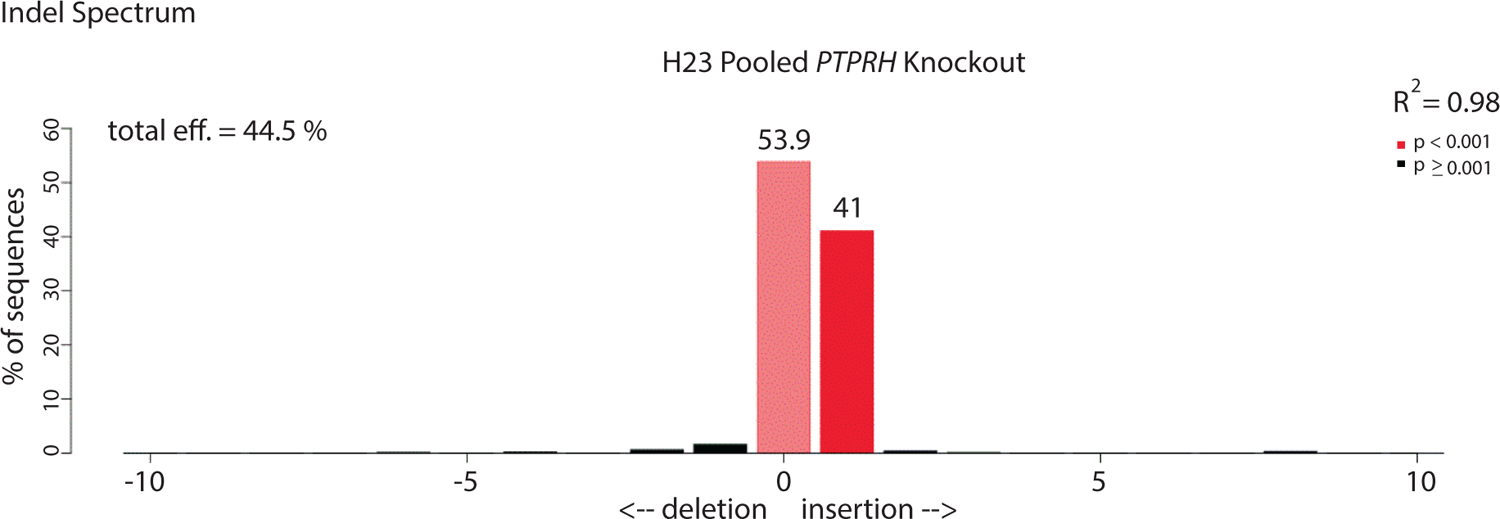

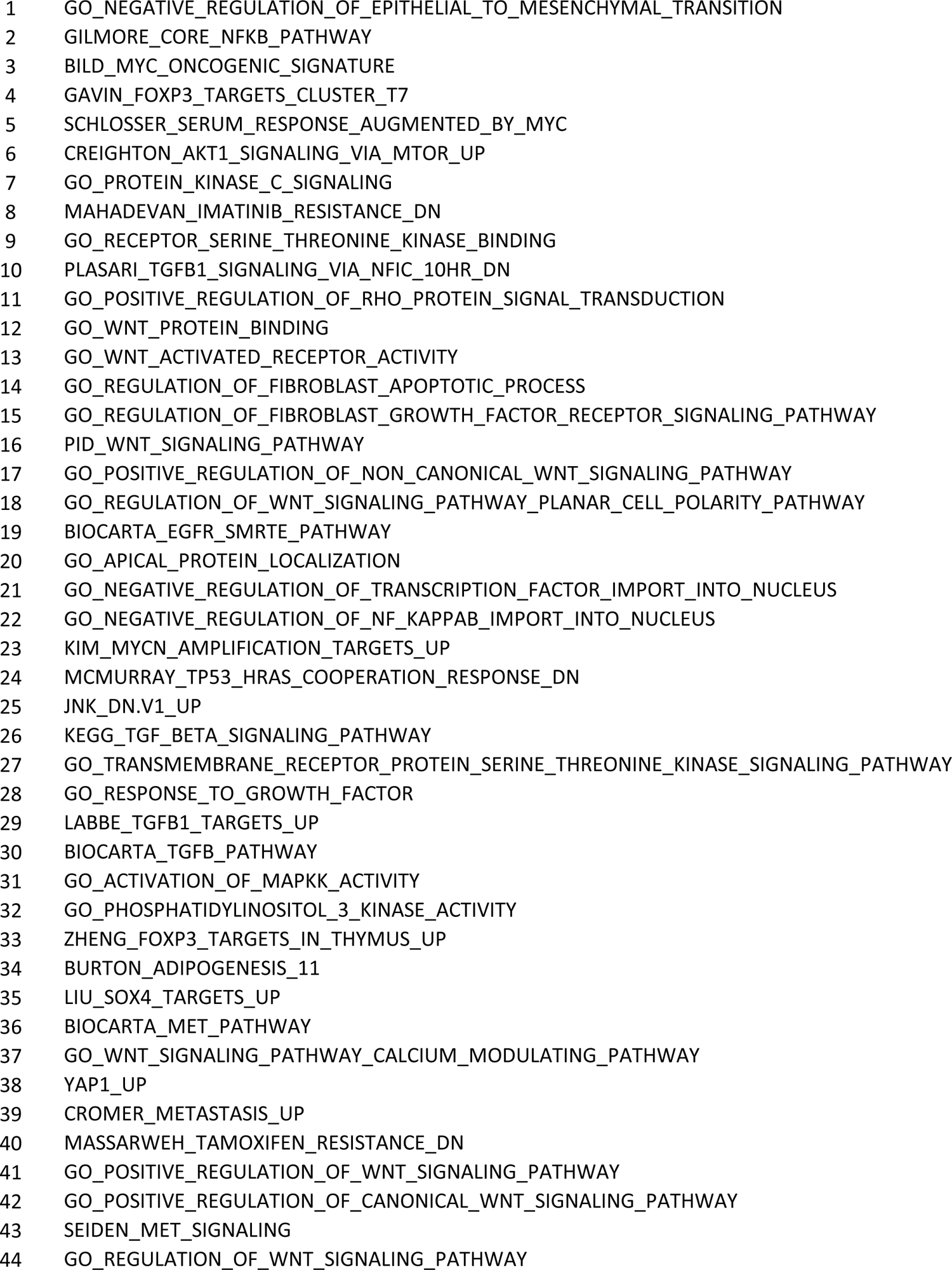
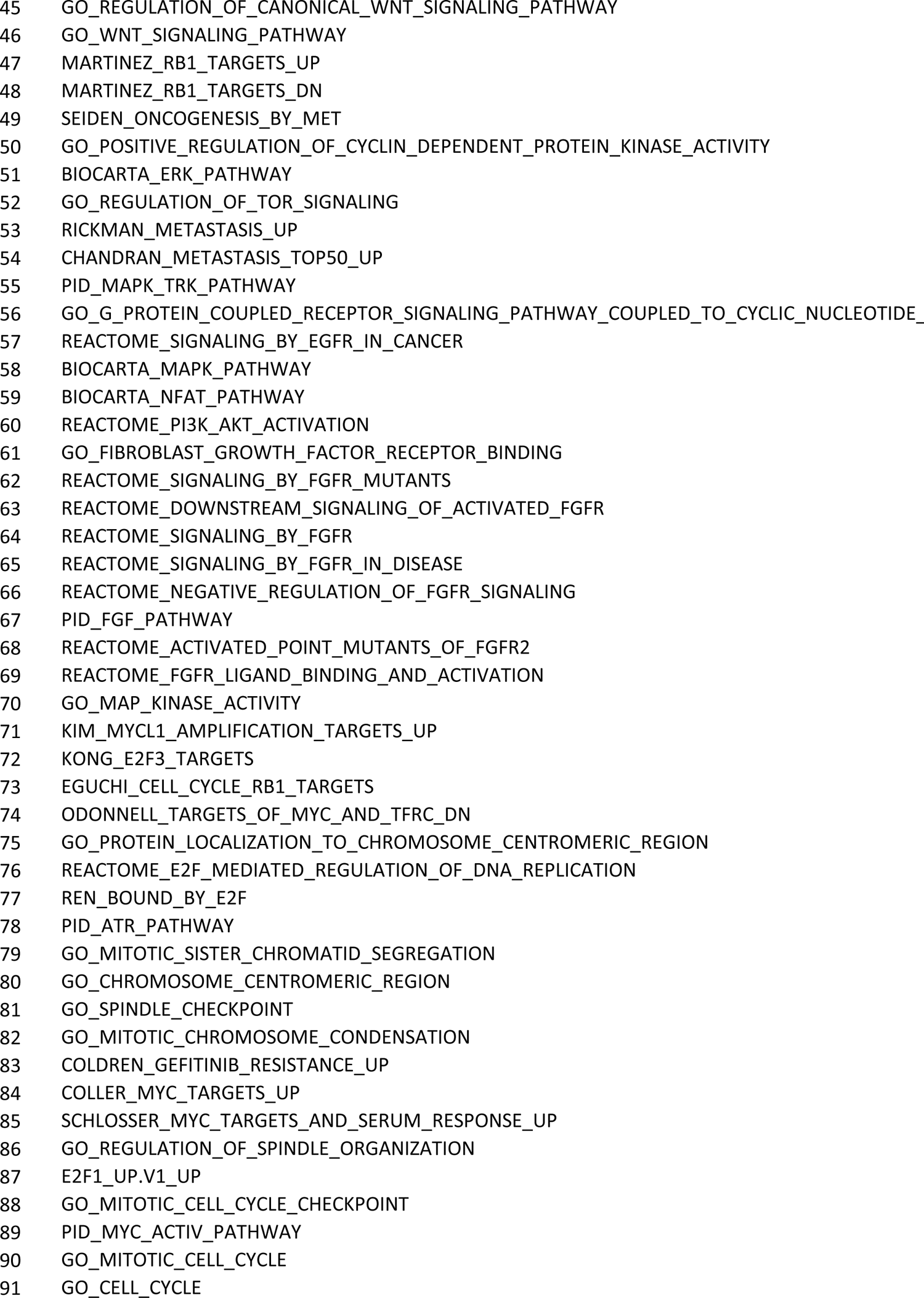
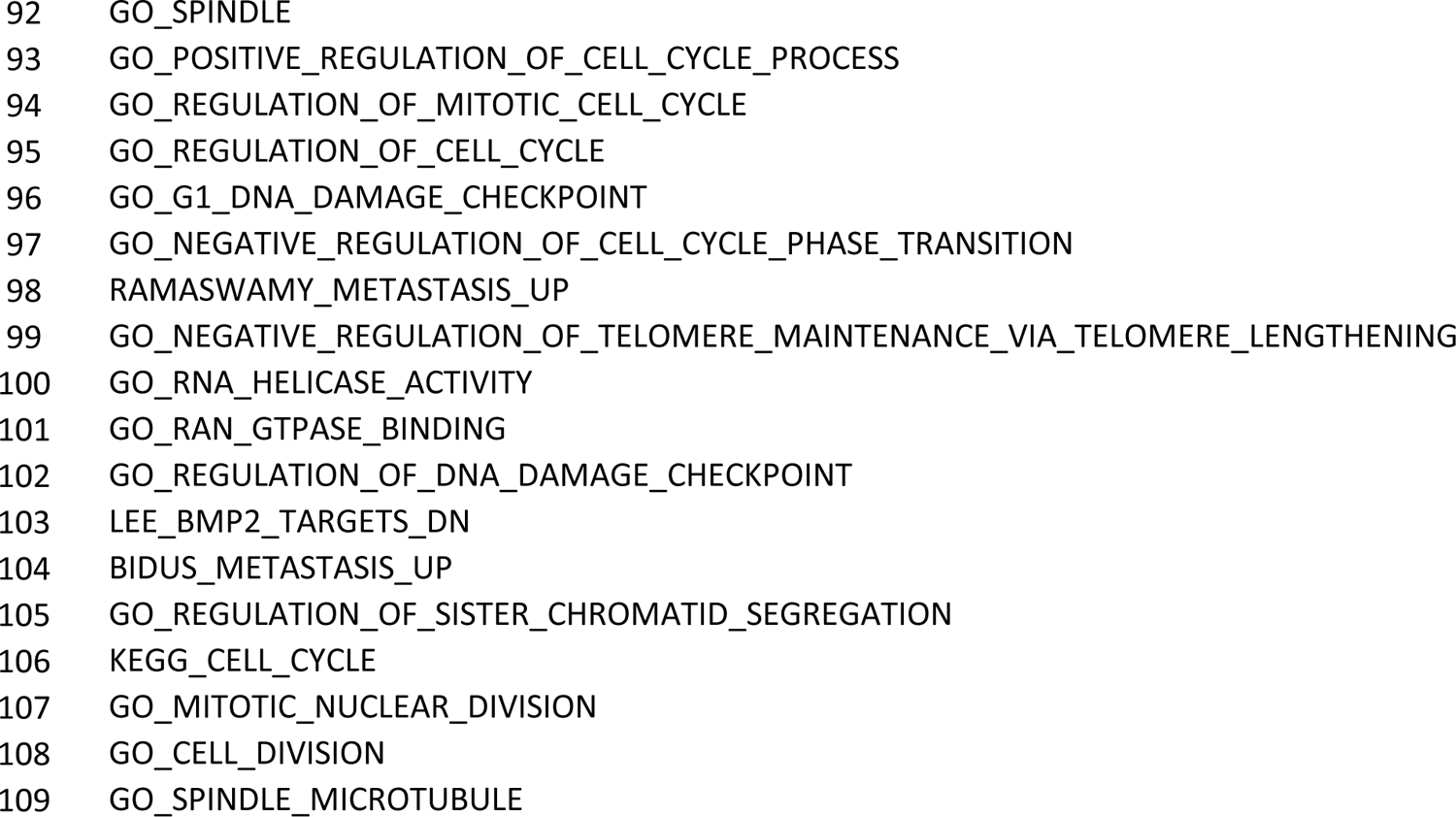

